# Dynamic Face-related Eye Movement Representations in the Human Ventral Pathway

**DOI:** 10.1101/2025.01.10.631890

**Authors:** Zhongbin Su, Xiaolin Zhou, Stefan Pollmann, Lihui Wang

## Abstract

Multiple brain areas along the ventral pathway have been known to represent face images. Here, in a magnetoencephalography (MEG) experiment, we show dynamic representations of face-related eye movements in the ventral pathway in the absence of image perception. Participants followed a dot presented on a uniform background, the movement of which represented gaze tracks acquired previously during their free-viewing of face and house pictures. We found a dominant role of the ventral stream in representing face-related gaze tracks, starting from the orbitofrontal cortex (OFC) and anterior temporal lobe (ATL), and extending to the medial temporal and ventral occipitotemporal cortex. Our findings show that the ventral pathway represents the gaze tracks used to explore faces, by which top-down prediction of face category in OFC and ATL may guide, via the medial temporal cortex or directly, face perception in the ventral occipitotemporal cortex.

---

Ventral occipitotemporal cortex is well known for its role in high-level object perception^1–6^. In particular, the fusiform face area (FFA) is activated by face images^7^ and the parahippocampal place area (PPA) by house images^8^. In a recent study, however, it was shown that FFA and PPA exhibited distinct neural activation patterns to face- and house-related gaze tracks, elicited in the absence of face or house image perception^9^. In this study, participants followed a sequence of dots on a uniform background with eye movements, where the dot sequence replayed gaze tracks previously recorded during face or house viewing. The face- and house-related gaze tracks could be decoded by the activation patterns in the FFA and PPA, thus indicating category-specific representations of gaze tracks in areas that were known to be activated by the respective image categories. Furthermore, the category-selective activation patterns were more sensitive to self-generated gaze tracks than gaze tracks generated by other observers^9^, in line with known individual differences in looking at faces^10^.

Here, we asked what the function of these gaze-track representations in a high-level perceptual area such as the FFA might be. Multiple areas along the ventral pathway are dedicated to face processing. In addition to the already mentioned FFA, face-responsive patches along the rostrocaudal extent of the temporal lobes as well as in the orbitofrontal cortex have been found in both human and non-human primates^11–13^. Notably, all of these studies investigated the processing of face images, without recourse to eye movements. Recently, however, modulations of neural activity by eye movements have been reported in the orbitofrontal cortex, particularly involving face viewing^14,15^. Moreover, neuronal activity in the medial temporal lobes can be modulated by eye movements, and lesions in this area may lead to changes in eye movement patterns during active sampling of the environment^16,17^. Medial temporal structures also interact with PPA during scene exploration^18^. Taken together with the evidence of gaze-sequence representation in the FFA and PPA^9^, these findings may lead to the hypothesis that the ventral stream, from orbitofrontal cortex via the uncinate fascicle to anterior temporal cortex and, via medial temporal cortex or directly, further to ventral occipitotemporal cortex, might be involved in representing face-specific gaze tracks.

However, how might face-specific gaze track representations be processed along the ventral stream? A top-down hypothesis is that eye movement sequences, e.g., during face viewing, initially activate prefrontal cortex, creating a categorical prediction (in this example of the ‘face’ category) which is fed back via the ventral stream to guide recognition in posterior perceptual areas^19–21^. To test this hypothesis, we investigated the spatiotemporal gradient of the MEG signal changes during face-related (vs. house-related) gaze track following. We expected dynamic signal changes specific to face-related gaze tracks to occur along the ventral stream, from orbitofrontal via anterior temporal and medial temporal cortex to ventral occipitotemporal cortex. Considering the known interindividual differences in the eye movement patterns of faces^10^, we also expected that following self-generated gaze tracks would optimally stimulate face-specific neural representations^9^, leading to earlier and stronger MEG signal changes than other-generated gaze tracks.

Top-down processing in the prefrontal cortex has been particularly observed when there is a lack of rich and unambiguous visual information^20,22^, as is the case in our dot-following task. Specifically, the categorical prediction in the prefrontal cortex was activated by ambiguous visual information in early visual areas^20^. To test this constraint, we presented actual images of faces and houses in the MEG scanner as a control condition. In contrast to the dot-following task, here, we expected a dominance of feedforward processing along the ventral pathway.

## Results

The study consisted of a behavioral experiment and an MEG experiment, with a one-week interval between the two experiments. In the behavioral experiment (Fig. 1a), gaze tracks of all participants were recorded while they were looking at images of faces and houses (see Table S1 in Supplementary Information for gaze parameters). In the MEG experiment, participants followed dot sequences of their own (self-face, SF; self-house, SH) or another participant’s gaze tracks (other-face, OF; other-house, OH) with eye movements (Gaze Session, Fig. 1b). After the Gaze Session, participants took part in an Image Session where they viewed images of faces and houses while maintaining the eyes on a central fixation point (Fig. 1c). The behavioral performance is presented in *Behavioral results* of Supplementary Information.

**Figure 1.**
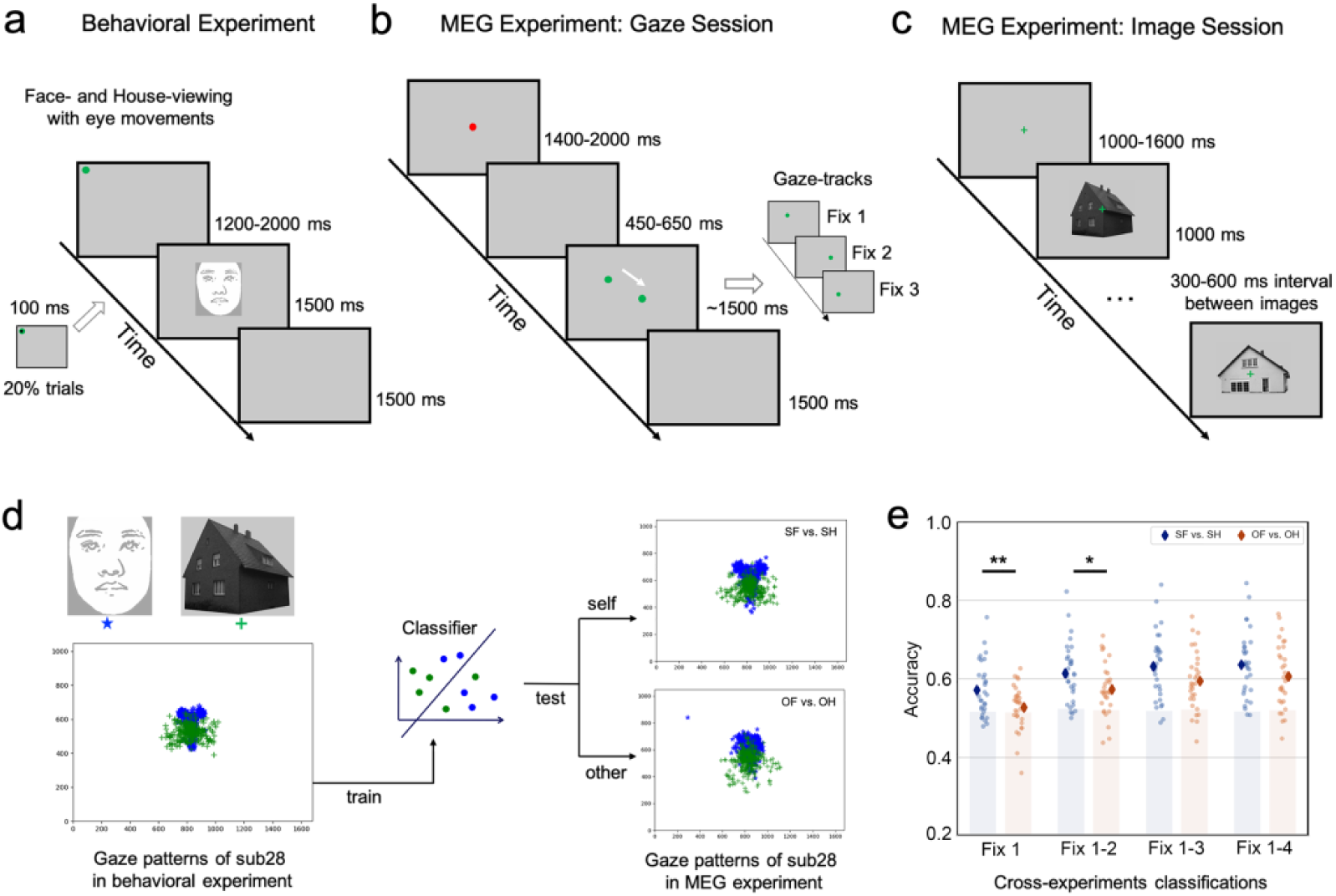
Experimental design and fixation decoding. (**a**) An example trial sequence in the behavioral experiment. (**b**) An example trial sequence in the Gaze Session of the MEG experiment. (**c**) An example block of the stimuli sequence in the Image Session of the MEG experiment. (**d**) An example face (blurred for privacy protection) and house image used in the experiment (upper left panel), and the fixation patterns (lower left panel) collected from one example participant during the view of faces (in blue star) and houses (in green cross). These fixation patterns were used to train a classifier in discriminating face- and house-related fixation patterns. The trained classifier was then used to classify the fixation patterns collected during the following of gaze tracks from the current observer (SF vs. SH, upper right panel) and the fixation patterns during the following of gaze tracks from another participant (OF vs. OH, lower right panel, i.e., cross-experiment classification). (**e**) The prediction accuracies of the cross-experiment classification are shown as a function of the comparison (SF vs. SH and OF vs. OH) and the number of fixations included in the classifications. The shaded areas indicate accuracies below the 95% percentile of the chance accuracies obtained from the permutation-based classification. \*\**p* < 0.01, \**p* < 0.05 for between self and other (Bonferroni-corrected).

### Distinct patterns between face- and house-related gaze tracks

We first analyzed if the face- and house-related fixation sequences obtained in the behavioral experiment showed distinct patterns. A machine-learning classification analysis over the spatiotemporal parameters of the fixations (*x*, *y* coordinates, and fixation duration) showed a high prediction accuracy in discriminating the two categories of gaze sequences, 70.5 ± 11.4% (M ± SD), with above-chance significance (*p* < 10^5^; permutation-based significance testing). The same analysis was also performed on the eye movements collected in the MEG-Gaze Session, yielding a high prediction accuracy in discriminating between SF and SH, 66.3 ± 10.2%, *p* < 10^5^, and between OF and OH, 66.5 ± 9.7%, *p* < 10^5^. We also performed cross-experiment classifications where the classifier was trained with the gaze patterns from the behavioral experiment and was used to predict the gaze patterns in the Gaze Session of the MEG experiment. The above-chance cross-experiment prediction accuracies confirmed that the distinct patterns of the online eye movement in the Gaze Session were related to the face vs. house categories in the behavioral experiment. Moreover, the cross-experiment prediction accuracies were higher for self-generated gaze tracks than for other-generated gaze tracks (Fig. 1e), indicating that participants followed their own gaze tracks better (see *Cross-experiment classification of gaze patterns* in Supplementary Information for the statistics).

### Neural face-related gaze pattern representations

To reveal the face-related gaze representations, we compared the MEG signals of face-related gaze tracks with the MEG signals of house-related gaze tracks. Here the house-related gaze tracks were taken as a control for face-related gaze tracks because there were eye movements and visual stimulation but a lack of a structural pattern^23^ (for statistical evidence, see *The structural pattern of face-related gaze trac*ks in Supplementary Information). For each condition (SF, SH, OF, OH) in the Gaze Session, the ERF signal from 0-1500 ms relative to the onset of the gaze track was calculated. The difference in the estimated cortical current maps was calculated between the following conditions: ‘SF – SH’, ‘OF – OH’, and ‘(SF – SH) – (OF – OH)’. These results would reveal the temporal development of the brain networks involved in the face-related gaze tracks, respectively areas that were sensitive to self-generated face-gaze tracks. The face-related gaze tracks elicited stronger ERF signals than the house-related gaze tracks in the orbitofrontal cortex (OFC) and the ventral anterior temporal lobe (ATL) extending to the medial temporal lobe (‘SF – SH’ and ‘OF – OH’ in Fig. 2a, Bonferroni-corrected for time and spatial clustering). Although small, there are also signal differences reaching back to the occipital cortex (at 600ms and 1400ms, depending on the contrast). Importantly, the activities in the OFC and ATL emerged earlier than the activities in the medial temporal lobe and occipital cortex, suggesting a top-down prediction of the face category guided by the face-related gaze tracks.

**Figure 2.**
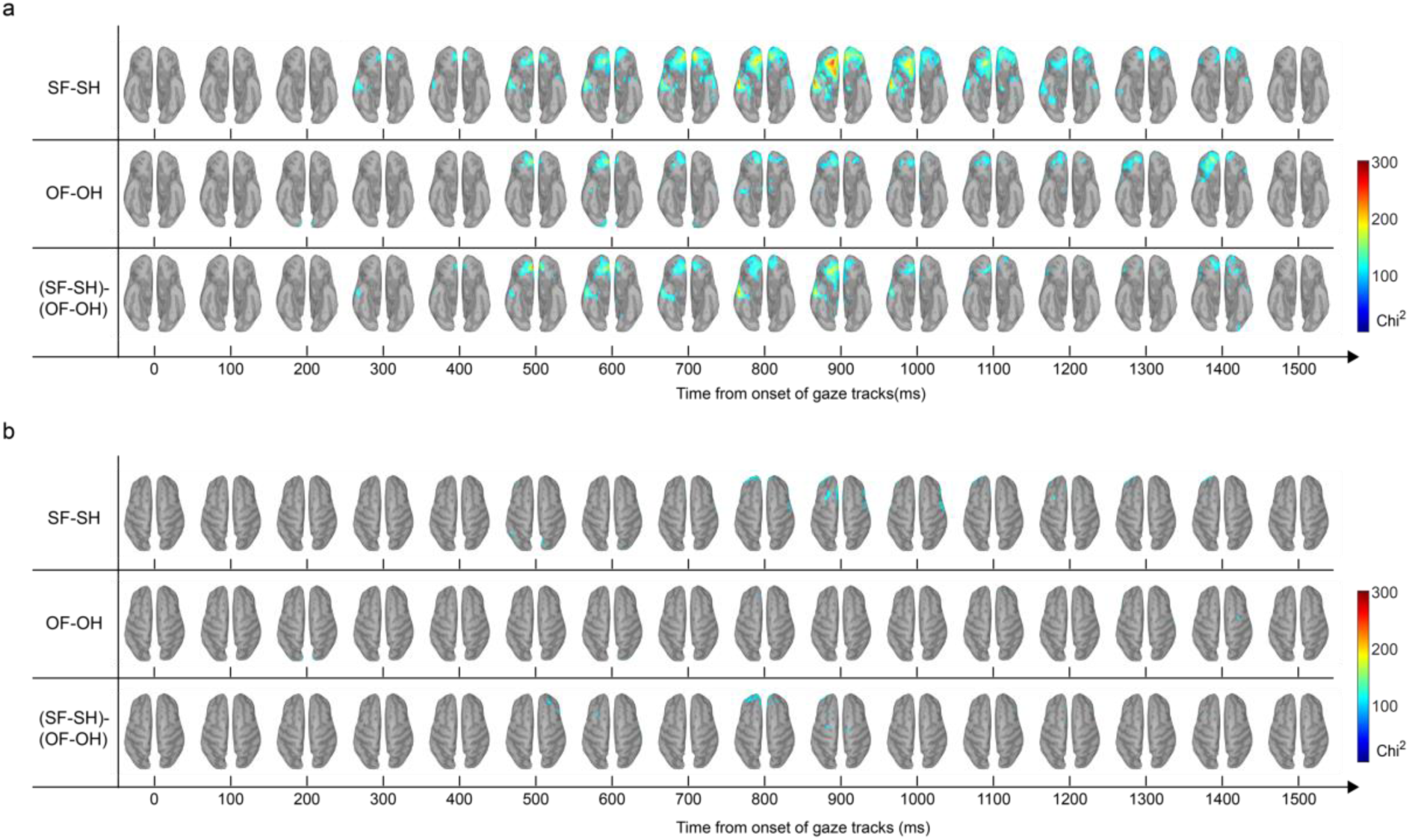
The ventral (**a**) and dorsal (**b**) view of the whole-brain source reconstruction based on the ERF signals. Upper row: the brain areas revealed by the contrast ‘Self-Face (SF) – Self-House (SH)’. Middle row: the brain areas revealed by the contrast ‘Other-Face (OF) – Self-House (OH)’. Lower row: the brain areas revealed by the interaction contrast ‘(SF – SH) – (OF – OH)’. The zero time points indicate the onsets of the gaze tracks.

Moreover, this network was more active during the following of self-generated gaze tracks than another observer’s gaze tracks, as revealed by the interaction contrast ‘(SF – SH) – (OF – OH)’ (Fig. 2a). The source reconstruction was dominantly localized in the ventral stream, with the notable exception of dorsal stream areas in frontal cortex, including the frontal eye field (FEF) and supplementary eye field (SEF), in the interaction contrast (Fig. 2b).

The whole-brain source reconstruction was also performed based on the signal difference between Face and House in the Image Session. In contrast to the Gaze Session, here the strongest signal difference was localized in the posterior occipitotemporal areas, beginning already 200 ms post-stimulus onset (Fig. 3).

**Figure 3.**
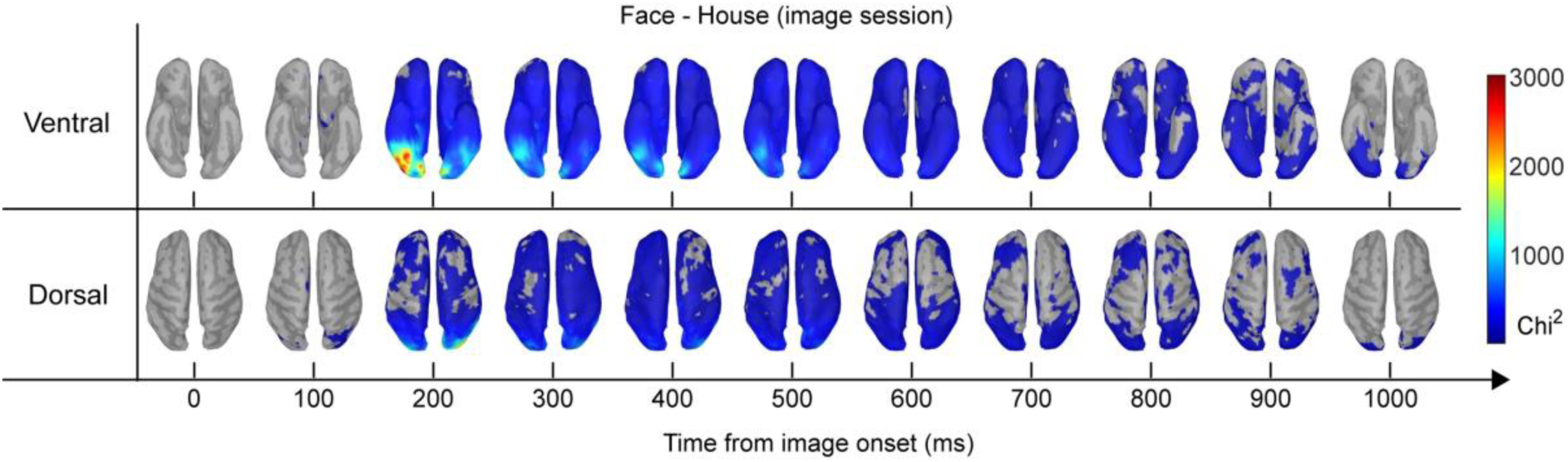
The ventral (upper row) and dorsal (lower row) view of the whole-brain source reconstruction based on the ERF signal difference between Face and House in the Image Session. The zero time point indicates the onset of the image.

### Dynamic spatiotemporal patterns of the gaze-related representations in the brain

In order to provide statistical evidence for the temporal order of signal development observed in the event-related magnetic field analysis, we tested if there was an information flow from the anterior areas (e.g., OFC and ATL) to the posterior areas (e.g., FFA) during gaze following. We performed spatial gradient analysis, which quantified how the MEG signals gradually changed along a specific dimension (e.g., anterior-posterior) in the brain space^24^. Here, the top-down hypothesis of the gaze-track representations predicted information flow from the anterior areas to the posterior areas along the ventral pathway. This can be probed with the gradually decreased activity from the anterior areas to the posterior areas, in particular, how the gradient became face-selective during the gaze following. As an area or neural network with stronger signals would be faster to exceed the neural threshold of maintaining sensory selectivity or perceptual preference^25^, the stronger signal changes in the anterior areas indicated that the face-selective representation of the gaze tracks emerged earlier than the posterior areas.

To test our hypothesis, here the spatial gradient was analyzed based on the ERF signal differences (e.g., ‘SF – SH’, ‘OF – OH’) to show how the signal changed at the anterior-posterior dimension. The analysis was performed at each time point during gaze following to show how the gradient pattern became face-selective over time. To provide a complete gradient pattern at the whole-brain level, the analysis was also performed for the dorsal-ventral and left-right dimensions. For each of the three dimensions (*x*, *y*, *z*, hence left-right, anterior-posterior, dorsal-ventral dimension), we modeled the coordinates with the ERF difference at each time point^24^. The *R*^2^ of the model was calculated to assess the accounted variance. The first-order derivatives of the estimated model were calculated to test if the spatial gradient increased or decreased monotonically along a specific dimension. As shown in Fig. 4a (left), the ERF signal difference between SF and SH showed a significant gradient pattern along the *y* and *z* dimensions, cluster-based permutation correction at *p* < 0.05 (see Fig. 4a for the significant time ranges), whereas the *x* dimension did not reach significance (no time ranges reached significance). The significant gradient pattern that emerged over time during the gaze following (i.e., significantly higher *R*^2^ than baseline) indicated that the gradient pattern was not due to the general intrinsic brain dynamics but rather to the face-specific gaze following. Along both the *y* and *z* dimensions, the estimated model showed a monotonic characteristic, with 97.5% of the derivative values > 0 at the time point with the strongest gradient pattern along the *y* dimension and 98.9% of the derivative values < 0 along the z dimension (Supplementary Fig. S2). The signal difference between OF and OH showed a similar pattern, with 98.7% of the derivative values > 0 along the *y* dimension and 97.7% of the derivative values < 0 along the *z* dimension (Fig. 4a right, Supplementary Fig. S2). These results indicated that the signal difference between SF and SH, and between OF and OH, decreased along the anterior-to-posterior axis (Fig. 4b upper row), and decreased along the ventral-to-dorsal axis of the brain (Fig. 4b lower row), whereas there was no difference between the left and right hemispheres. Taking the results at the anterior-posterior and the ventral-dorsal dimensions together, these findings provided statistical evidence for the temporal sequence observed in the event-related magnetic field analysis, showing that the representations of face-related gaze tracks dynamically progressed along the ventral pathway, starting from the ventral anterior areas (e.g. OFC and ATL) via medial temporal lobe (MTL) to ventral occipitotemporal cortex.

**Figure 4.**
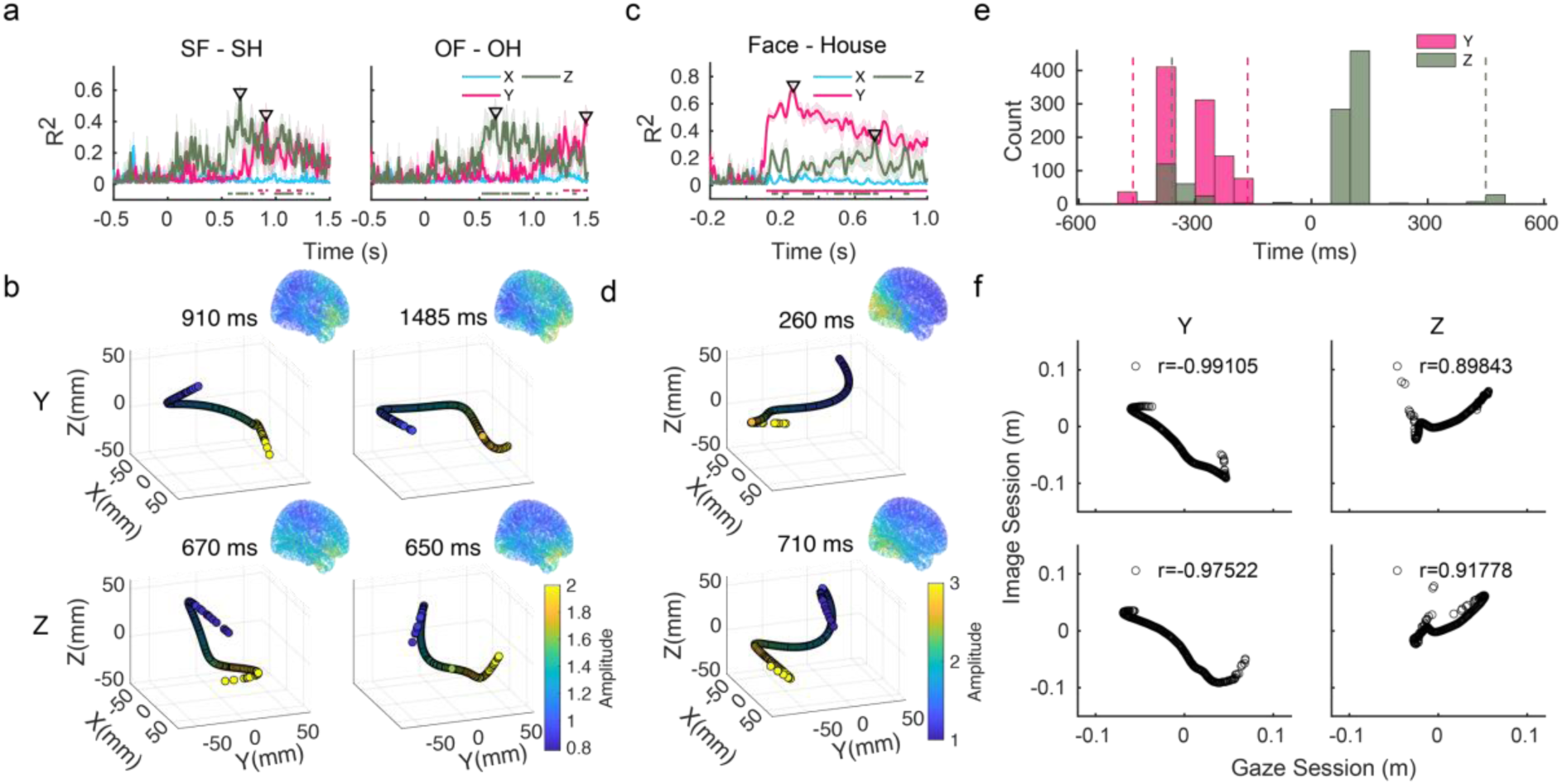
The spatial gradient of the MEG signal at each dimension of the 3-D brain space. The *R*^2^ of the spatial gradient model are shown as a function of the spatial dimensions (*x*, *y*, *z*) and time in the Gaze Session (**a**) and the Image Session (**c**). The horizontal lines at the bottom of each pallet indicate the time ranges where the *R*^2^ were significantly higher than the baseline (multiple comparisons corrected with cluster-based permutation at *p* < 0.05). The small triangles indicate the peak of the *R*^2^ along a specific dimension. The spatial gradient patterns (in terms of *R*^2^ values) at the peak time points are shown in the 3D space (**b**: Gaze Session, **d**: Image Session). (**e**): The counts that the spatial gradient pattern for ‘SF – SH’ emerged earlier than the spatial gradient pattern for ‘OF – OH’ along the *y* and the *z* dimensions. The latency difference between the two *R*^2^ time courses was estimated with a cross-correlation method and was tested using a bootstrapping method (see Methods). Dashed lines indicate the 95% confidence interval. (**f**): The predicted coordinates sorted by the ERF amplitudes in the Image Session are shown as a function of the predicted coordinates in the Gaze Session along the *y* (left column) and the *z* (right column) dimensions. Upper row for ‘SF – SH’ and lower row for ‘OF – OH’.

The spatial patterns at the time point where the gradient reached its peak are shown in Fig. 4b (left: ‘SF – SH’, *R*^2^ peaked at 910 ms along the *y* dimension and at 670 ms along the *z* dimension; right: ‘OF – OH’, *R*^2^ peaked at 1485 ms along the *y* dimension and at 650 ms along the *z* dimension). Importantly, the spatial gradient emerged earlier for ‘SF – SH’ than the spatial gradient for ‘OF – OH’ along the *y* dimension (i.e., the posterior-to-anterior dimension), mean latency difference = −310 ms, 95% CI = [−460 ms, −165 ms], but not along the *z* dimension mean latency difference = 20 ms, 95% CI = [-360 ms, 450 ms] (Fig. 4e). Together with the cross-experiment classifications of gaze patterns, these results indicated that self-generated gaze tracks were more sensitive than other-generated gaze tracks to activate the face-selective neural representations.

In the Image Session, the ERF signal difference between Face and House also showed a significant gradient pattern along the *y* and *z* dimensions, cluster-based permutation at *p* < 0.05, whereas the *x* dimension did not reach significance (Fig. 4c). Along both the *y* and *z* dimensions, the estimated model showed a monotonic characteristic, with 99.4% of the derivative values < 0 along the *y* dimension and 82.3% of the derivative values < 0 along the *z* dimension (Supplementary Fig. S2). The spatial patterns at the time point where the gradient reached its peak are shown in Fig. 4d (*R*^2^ peaked at 260 ms along the *y* dimension and at 710 ms along the *z* dimension). While the gradient pattern along the *z* dimension was consistent with the Gaze Session (Fig. 4c), with signal difference decreasing from the ventral to the dorsal part of the brain (Fig. 4d, lower row), the gradient pattern along the *y* dimension was reversed, with signal difference decreased from the posterior to the anterior part of the brain (Fig. 4d, upper row). Importantly, the reversed gradient pattern along the *y* dimension between the Gaze Session and the Image Session again indicated that the observed gradient pattern was not due to the general intrinsic brain dynamics along the ventral pathway, but rather reflected the feedback-dominant vs. feedforward-dominant processing specific to the current task (i.e., gaze following vs. image processing).

The reversed pattern along the anterior-posterior direction between the Gaze Session and the Image Session was further confirmed by the statistical evidence that the spatial gradient along the *y* dimension showed a negative correlation between the two sessions, *r* = −0.99, 95%CI = [−0.994, −0.957], *p* < 0.001 at the peak time point for ‘SF – SH’, and *r* = −0.98, 95%CI = [−0.991, −0.934], *p* < 0.001 at the peak time point for ‘OF – OH’ (Fig. 4f, left). By contrast, the spatial gradient along the *z* dimension showed a positive correlation between the two sessions, *r* = 0.90, 95%CI = [0.863, 0.951], *p* < 0.001 at the peak time point for ‘SF – SH’, and *r* = 0.92, 95%CI = [0.856, 0.953], *p* < 0.001 at the peak time point for ‘OF – OH’ (Fig. 4f, right). Collectively, the reversed pattern along the anterior-posterior direction between the Gaze Session and the Image Session suggested a combination of feedback and feedforward processing in natural face perception (i.e., when we look at a face using eye movements).

## Discussion

We have shown that face-related gaze tracks were dynamically represented along the ventral pathway, from OFC via ATL, to MTL and ventral occipitotemporal cortex. During the gaze following, there was a gradient pattern along the ventral stream, with face-selective activity progressing from OFC to the occipitotemporal cortex. However, when actual images of faces and houses were presented, the reverse gradient was observed, face-selective activity progressing from the ventral posterior occipitotemporal cortex to the prefrontal cortex. Taken together, our findings show that the ventral pathway represents aspects of the eye-movement program used to explore faces. The fixation sequences may help to form a top-down prediction of face category in OFC and ATL to guide, via the MTL or directly, the perceptual representation of faces in ventral occipitotemporal cortex, particularly under demanding viewing conditions.

The brain areas that we found representing categorical face-related gaze tracks, from OFC via ATL to ventral occipitotemporal cortex have previously been found to support face perception, both in human and non-human primates^11–13^. Importantly, we found these areas to represent face-specific gaze patterns in the absence of face or house images, indicating that not only visual features, but also information about category-specific gaze sequences are represented, like fixation locations and their temporal sequence. In addition, the representation of face-related gaze sequences was particularly early and strong when gaze sequences that were followed were generated by the same participant during actual face viewing. This pattern is in line with the stable interindividual differences in gaze patterns that can be found across different viewing conditions^10^.

Face-specific activity was observed earliest in OFC and ATL, spreading backwards along the ventral stream. Although we do not know about previous reports of eye movement processing in human orbitofrontal cortex, modulation of orbitofrontal activity by eye movements has recently been reported, particularly during looking at faces^14,15^. OFC and ATL, connected via the uncinate fasciculus, are known to be vital for social interaction, with lesions in this network leading to the behavioral variant of frontotemporal dementia^26,27^. Given the importance of face perception - including perception of facial expressions - for social interaction, it is not astonishing to find representations of face-specific gaze sequences in these areas. The early occurrence of face-specific gaze representation in OFC and ATL in the ventral face processing stream may suggest that the processing of face-specific gaze patterns is vital for social interaction. It may thus be worthwhile to investigate if face-specific gaze sequences break down in degenerative diseases affecting OFC and ATL, like frontotemporal dementia.

Face-related gaze sequence activation spread further to the medial temporal lobe, which has previously been shown to be vital for the exploration of the environment with eye movements, particularly, but not only, in memory-guided vision^28–30^. Moreover, during face viewing, activation in the hippocampal as well as in the fusiform face area was modulated by the number of fixations^31^ and during scene viewing, functional connectivity between the hippocampus and the PPA was enhanced during free viewing (versus forced central fixation)^18^. The MTL is connected to orbitofrontal and temporopolar cortex via the perirhinal cortex^32^. Thus, in the context of face viewing, information from OFC and ATL about the highly structured (T-shaped) fixation pattern may elicit a memory trace of the ‘face’-category in the hippocampus. This, however, needs further investigation.

If OFC and ATL are first to represent face-specific gaze patterns, how does the information about fixation patterns arrive at these areas in the first place? In normal looking behaviour, this may be answered by our finding that during the presentation of actual face images, faces and houses can be discriminated early and most strongly in occipitotemporal cortex, spreading fast to anterior brain areas. Thus, during free viewing of a face, there will be an interaction of feedforward and feedback signals supporting perception. Our data, in line with previous reports, suggest that information about face-specific gaze sequences may aid face perception particularly in the absence of rich and unambiguous visual information^19,20,22^. A recent MEG study also showed that the ventral prefrontal cortex guides the construction of low-dimensional categorical prediction from the high-dimensional visual information in occipitotemporal cortex^21^. Taken together, following the gaze tracks may activate category-predictive processes in OFC and ATL, sending feedback signals via the MTL or directly to ventral occipitotemporal face patches including the FFA to facilitate face perception.

Recently, it has been emphasized that categorical object representation in the brain may be linked to the object’s behavioral relevance^33^, in line with models of perception-action coupling^34–36^. Given that eye movements are mostly generated fast and without conscious control^37^, representations of category-specific gaze sequences as part of an object’s neural representation may be a natural case of object representation including relevant behavior, at least for object categories with structured gaze sequences, such as for faces.

It may seem puzzling that we found gaze-specific activation patterns for faces mainly in areas of the ventral stream, whereas the dorsal stream’s importance for eye movement control is well known^38–41^. We also could discriminate face versus house-related gaze following in dorsal brain areas, particularly in the frontal eye fields and the superior frontal cortex, known to support attentional control^42^, as well as in left frontopolar cortex, known to support exploratory attentional resource allocation^43^. Interestingly, this was only the case for self-generated dot following, ruling out that these activation patterns were simply due to differences in basic eye movement parameters like saccade amplitudes. Nevertheless, dorsal stream activation differences for face versus house-related gaze following were much less than in ventral stream areas. This may be due to the nature of our contrasts, which asked for a categorical (‘face – house’) distinction between the associated gaze sequences. This categorical distinction may be more associated with the known capabilities of the ventral stream in object categorization than with the visuomotor control functions associated with the dorsal stream^44^.

## Conclusion

A ventral network of brain areas known to support face processing, reaching from orbitofrontal and anterior temporal cortex via medial temporal to ventral occipitotemporal cortex, was found to represent face-related gaze sequences. During gaze following, activation in this network followed an anterior-to-posterior gradient, indicating feedback from anterior areas to ventral occipitotemporal perceptual areas, possibly using gaze patterns to support face perception. Our findings add an ideomotor perspective to the ventral stream function and the reconsideration of both ventral and dorsal streams in representing eye movements. The face-related eye movement representations in OFC and ATL suggest the role of the frontotemporal network in gaze control during social interaction.

## Methods

### Participants

The sample size was decided with G-Power 3.0^45^ based on a previous study that examined the capabilities of MEG signals in decoding eye movement patterns^46^. Given a correlation coefficient between eye movement and MEG patterns = 0.86, alpha = 0.0001 reported in this study, a sample size of 27 is required to achieve an expected power of 99%. Following this criterion while considering potential exclusion, 32 university students (19 females, 13 males, mean age 20.8 years old) were recruited in the present study. All participants reported normal or corrected to normal vision. Informed written consent was obtained prior to the experiment. One participant was excluded due to his drop-out from the MEG experiment, resulting in 31 participants (19 females, 12 males, mean age 20.6 years old). This study was conducted in accordance with the Declaration of Helsinki, and was approved by the ethics committee of the local university.

### Design and procedure

Each participant went through a behavioral experiment (Figure 1a) and an MEG experiment, with a one-week interval between the two experiments. In the behavioral experiment, we collected the gaze patterns of all participants while they were looking at faces and houses. These gaze patterns were then presented in the Gaze Session of the MEG experiment, and participants had to follow the gaze patterns with eye movements.

In the behavioral experiment, stimuli were images of faces and houses presented on a black background of a computer screen. For each participant, the images of faces and houses presented during the experiment were randomly chosen from an image set (20 male faces, 20 female faces, and 40 houses). The size of the face images was fixed at a width of 14.4° × height of 16.7° of visual angle, with an eye-to-mouth distance of 7°. Due to the varying structures, the size of houses was not constant, with a mean width of 19.3° ± 1.0°, and a height of 13.1° ± 2.4°. Participants were required to complete an N-back task while looking at each of the pictures^9^. At the beginning of each trial, a green dot (0.2° of visual angle in diameter) was presented at one of the four corners (15° from the center of the screen) to attract eye fixation. The green dot was presented for a varying interval of 1200-2000 ms. In 20% of the trials, a small black dot (0.05° in diameter) was presented at the center of the green dot for 100 ms. Participants were asked to detect the black dot by pressing the ‘z’ button using the left index finger on a standard keyboard. The onset of this small black dot was randomly chosen from the time point during the presentation of the green dot. After the offset of the green dot, a face or house picture was presented at the center of the screen and remained on the screen for 1500 ms. Participants were instructed to look at the images with free eye movements. There were 14 blocks of trials in the experiment. In the first block, a set of 6 face images (3 males and 3 females) and 6 house images were presented, one per trial, in a random order. In each of the following 13 blocks, 1-3 new images (either face or house) were added into the original 12 images. Participants were asked to memorize the images in the first block, and detect if a new image was presented in the following 13 blocks by pressing the ‘m’ button using the right index finger.

In the MEG experiment, stimuli were presented through an LCD projector onto a rear screen located in front of the scanner. There were two sessions in the MEG experiment: a Gaze Session and an Image Session.

In the Gaze Session, each participant was asked to follow dots that represented his/her own gaze patterns as well as dots that represented another participant’s gaze patterns. Each trial started with a red dot on a grey background, which remained at the center of the screen for 1400-2000ms. After a blank screen of a jittered interval (450 – 650 ms), the gaze track was presented on the screen, in the form of a sequence of green dots. The sequence of green dots represented the gaze pattern for a specific picture obtained from the behavioral experiment, and each dot represented a fixation of the gaze pattern. Given that the gaze patterns were collected during the 1500ms-time range of the picture presentation, the dot sequence lasted approximately 1500 ms on the screen. In 20% of the trials, a small black dot (0.05° in diameter) was presented at the center of the central red dot or the moving green dot (with equal probabilities). This small black dot was presented for 100 ms, and participants were asked to detect the black dot by pressing the button using the right index finger. According to our design, four categories of gaze tracks were presented: face-related gaze tracks from the current observer (self-face, SF), house-related gaze tracks from the current observer (self-house, SH), face-related gaze tracks from another participant (other-face, OF), and house-related gaze tracks from another participant (other-house, OH). There were 10 blocks of gaze-tracks, with 40 trials (10 trials per condition) in each block. Trials from the 4 conditions were mixed and presented in a random order. At the end of each block, a feedback screen was presented to inform the participants’ performance in detecting the small black dots.

In the Image Session, participants were asked to view successively presented images in a one-back task. Images were grouped into 10 blocks of faces and 10 blocks of houses. The two block types were presented in a random order. Each block started with a central fixation (a green cross) at a varying interval of 1000-1600 ms. Then images of the same category (face or house) were successively presented (each lasted for 1000 ms), with a jittered interval of 300-600 ms between each two images. The central fixation was presented throughout the block, and participants were required to maintain their eyes on the central fixation. In each block, 10-11 images were presented with one or two images that were repeated immediately after their first presentation. Participants were asked to detect the immediate repetition of the image by button press. Apart from the immediate repetition, there were no other repetitions of images in each block. In total, each participant viewed 100-110 images of each category. There was a ∼30s break between each two blocks.

For both the behavioral experiment and the MEG experiment, eye-movement data were recorded during the experiment with an EyeLink 1000 plus system (SR-Research, Canada), at an online sampling rate of 1000 Hz. A standard procedure of nine-point calibration and validation was performed at the beginning of the experiment, with a maximum error of 1.0° as the threshold. A drift check was performed at the beginning of each block, and the calibration and validation were performed if the error of the drift check exceeded the threshold (i.e., > 1.0°).

### MEG data acquisition and preprocessing

Neuromagnetic signals were recorded using a whole-head MEG system, with 204 planar gradiometers and 102 magnetometers (Eleka Neuromag TRIUX) in a magnetically shielded room. Four head position indication (HPI) coils were placed in each participant’s head to estimate head position during recording, with two coils in left and right mastoids and two on the forehead. The raw MEG signals were online sampled at 1000Hz and were band-pass filtered between 0.1 and 330 Hz. The structural MRI of each participant was obtained using a 3T Siemens Prisma MR scanner. The MRI scanning was conducted on a different day after the MEG experiment.

Head shapes were quantified using the Probe Position Identification system (Polhemus), and three anatomical landmarks (nasion, left and right pre-auricular points) were used to co-register the MEG data with MRI coordinates. Max-filter was used to reduce external noise and compensate for head movements (temporal signal space separation method, tSSS^47^). The offline pre-processing analysis of MEG data was performed using Brainstorm^48^. The continuous MEG data was first down-sampled to 200 Hz. Then the MEG data was band-pass filtered (0.1Hz to 60Hz, zero phase shift FIR filter) and notch filtered at 50 Hz. Independent component analysis (ICA) was used to detect and discard artifacts related to eye blinks, head movements and heat beats. The data were then epoched with the time interval of −500 to 1500 ms relative to the onset of the first fixation in the Gaze Session and with the time interval of −200 to 1000 ms relative to the onset of the image in the Image Session.

### Analysis of eye-movement data

In the behavioral experiment, eye-movement data were extracted from the 1.5-s image presentation. Data were preprocessed using the *cili* module, a python-based tool for detecting and correcting eye blinks. Eye blinks were firstly removed, and fixations were identified based on the velocity threshold of 30 °/s and the acceleration threshold of 8000°/s^2^. Trials without any valid fixation events, and trials with fixation localized beyond the region of the picture were also excluded. To prepare the gaze tracks in the MEG experiment, following the previous study^9^, a fixation was identified as a gaze event if its duration was longer than or equal to 100 ms, while identified as a non-gaze event if its duration was shorter than 100 ms. This non-gaze event was represented by a blank screen in the Gaze Session of the MEG experiment. Then, trials with less than two gazes were excluded. The gaze coordinates were proportionally transformed and co-registered with the screen resolution in the MEG scanner. In the Gaze Session of the MEG experiment, the online fixation events were also identified and analyzed.

Multivariate classifications were performed on the gaze features to show the distinct patterns between categories. The classification analysis was performed using the scikit-learn package (http://github.com/scikit-learn). Three features were included: the *x*, *y* coordinates, and the duration of each gaze. The fixation data was parsed in the way that 80% of the data was included as the training set and 20% of the data as the test set. A linear support vector (SVM) classifier was trained and cross-validated based on the saccadic features of the two categories (Face vs. House). The classification was performed for each participant, rendering both individual-level prediction accuracies and the group mean of the accuracies. Permutation-based testing was conducted to assess the statistical significance. For each participant, the classifier was trained with randomly shuffled labels of the two categories, and a permuted accuracy was calculated. This procedure was repeated 100 times, rendering a set of 100 chance accuracies for each participant. For group-level statistical testing, one chance accuracy was selected from each participant and the individual chance accuracies were averaged into a group chance accuracy. This procedure was repeated 10^5^ times, resulting in a set of 10^5^ group chance accuracies. Significance testing was performed by calculating the probability of the unpermuted group mean accuracy across participants in the distribution of the group chance accuracies (one-tailed). The classification was performed both for the fixation data in the behavioral experiment (Face vs. House) and the fixation data in the Gaze Session of the MEG experiment (SF vs. SH, OF vs. OH).

Note that the SF vs. SH and OF vs. OH classifications in the Gaze Session were not strictly specific to the Face vs. House distinction because no face or house images were presented. To show the specificity, cross-experiment classifications were performed where the fixation patterns in the behavioral experiment were used to train the classifier (Face vs. House), which was then used to predict the fixation categories in the Gaze Session (SF vs. SH, OF vs. OH). Importantly, to assess the sensitivity of the distinct fixation patterns, the cross-experiment classification was performed by varying the number of the fixations (i.e., the first fixation, the 1-2 fixations, the 1-3 fixations, and the 1-4 fixations). Multiple comparisons were corrected with Bonferroni methods.

### Event-related magnetic field (ERF) analysis of MEG data

After the pre-processing, the epoched data were averaged over the trials for each condition and each participant. Individual T1-weighted MRIs were segmented with the FreeSurfer software package^49^ (http://surfer.nmr.mgh.harvard.edu) and then imported to the Brainstorm (https://neuroimage.usc.edu/brainstorm) for further source-level analysis. The white-gray matter boundary segmented by the FreeSurfer was used as a source space for activity estimation in the cortex. After co-registration between the individual anatomy and MEG sensors, the cortical currents were estimated using a distributed model consisting of 15002 current dipoles from the averaged epochs (evoked activities) using a linear inverse estimator (minimum norm current estimation). The density map was standardized using a Z-score transformation with respect to a noise matrix which was calculated with a 2-minute empty-room recording of the MEG signal. The dipole orientation was constrained to the orthogonality of the white-gray matter boundary of the individual MRIs.

The difference in the estimated cortical current maps was calculated between the following conditions: ‘SF – SH’, ‘OF – OH’, ‘(SF – SH) – (OF – OH)’. Then the source maps were filtered with a low-pass filter (30Hz), standardized through a z-score baseline normalization (−450 to 0 ms relative to the gaze onset as the baseline, with the first 50 ms of the baseline period being excluded to avoid the edge effect resulted from the low-pass filter), and rectified to retain only absolute values. The source maps were then projected on a standard brain (ICBM152) and spatially smoothed (Full Width at Half Maximum, FWHM=3mm) before group statistical analysis. A two-tailed one-sample Chi^2^ test was used for group statistical analysis for each time point and each vertex with the null hypothesis that the difference in variances of the cortical activities between the two conditions was equal to zero^50^. Bonferroni correction was used to solve the multiple comparison problems. The significance threshold was set at *p* < 0.05 after corrections.

To show the brain areas that were involved in the Image Session, the whole-brain source reconstruction was also performed by comparing the ERF signals during face viewing and the ERF signals during house viewing (‘Face – House’).

### Modeling the gradient of MEG signal in source space

To quantify the spatial patterns of MEG signals during the gaze-track following, a five-order polynomial function was used to approximate the data along each of the three spatial dimensions (*x*, *y* and *z* coordinates for the data in the source space)^24^. For each time point, we employed a polynomial function *p(v) = p_0_ + p_1_v + p_2_v^2^ + p_3_v^3^ + p_4_v^4^ + p_5_v^5^*(*polyfit*, MATLAB 2022a) to estimate the coordinates along each spatial dimension with the MEG signal difference between conditions (e.g., ‘SF – SH’, ‘OF – OH’). The amplitude of MEG signal was normalized to z-scores across vertexes to avoid an ill-conditioned Vandermonde matrix in model fitting. For each spatial dimension, the model fitted the signal difference in Z-scored amplitudes of each vertex *v* to its spatial coordinate (MNI coordinates) across vertices. The quality of the model was quantified by the adjusted *R*^2^, which determined the proportion of variance explained by the model. *R*^2^ was adjusted by the number of coefficients. To assess the dynamic spatial gradient of the MEG signal difference, a Jackknife method was used to fit the model and calculate *R*^2^ for each of the 3 dimensions. Specifically, one of the participants was excluded and the source-reconstructed MEG signals of the remaining participants were averaged to fit the model. This procedure was iterated across participants. A one-sample *t* test (one-tail) was used to test if *R*^2^ at each time point was higher than the baseline, the time interval of −500 to 0 ms relative to the stimulus onset. Cluster-based permutation was used to resolve the multi-comparison problem across time points. We also calculated the first-order derivatives of the estimated model to test if the spatial gradient had a monotonic increasing or decreasing pattern along a specific dimension. The calculation was performed on the model at the time point with peak *R*^2^, and the evaluation of the derivatives was based on the signal range between the minimum and the maximum value of the MEG amplitude. The spatial gradient was identified as monotonically increasing given the derivative values > 0 and monotonically decreasing given the derivative values < 0.

To test if the spatial gradient of ‘SF – SH’ emerged earlier than the spatial gradient of ‘OF – OH’, cross correlation (*xcorr*, MTALAB 2022a, ‘unbiased’, maxlag = 200) was performed on the two *R*^2^ time courses to calculate the latency difference. The latency difference was defined as the temporal lag with which the *R*^2^ time courses showed maximum correlation between the two time courses across participants. The Bootstrapping method (iteration number = 1000) was used to estimate the 95% confidence interval of latency difference.

The same analysis was also performed on the MEG signal difference between Face and House in the Image Session to show the spatial gradient. The analysis was performed on the 0-1000 ms interval during the image presentation (0 denotes the image onset), with the - 200-0 ms interval as the baseline.

To assess the similarity/dissimilarity of the spatial gradient between the Gaze Session and the Image Session, we performed correlation analyses between the two sessions. Specifically, we performed the model fitting with the group average of source-reconstructed MEG signal difference between conditions (‘SF – SH’ and ‘OF – OH’ for the Gaze Session and ‘Face – House’ for the Image Session). A Bootstrapping method (iteration number = 1000) was used to estimate the variance of *R*^2^ time courses that were calculated between the model and the group average of the MEG data. At the peak time point of *R*^2^ time courses (*y* and *z*, respectively), we projected the fitted function *p*(v) in the three-dimensional space with the polynomial function. The predicted coordinates of the function *p*(v) were sorted according to the amplitude of the MEG signal. Then we calculated the *Pearson* coefficients of the sorted coordinates between the two sessions.

## Acknowledgments

We thank Dr. Jiayu Zhan and Dr. Liyu Cao for their suggestions on the design of the MEG experiment. This study was supported by the National Natural Science Foundation of China (32271086), and a Mercator Fellowship of the Deutsche Forschungsgemeinschaft (DFG, 450600965) to LW, and a DFG grant (PO548/18-1) to SP.

## Author contributions

Conceptualization, L.W., S.P., Z.S.; Methodology, Z.S., L.W.; Investigation, Z.S., L.W.; Formal Analysis, Z.S., L.W.; Visualization, Z.S.; Writing – Original Draft, S.P., L.W., Z.S.; Writing – Review & Editing, S.P., L.W., Z.S., X.Z.; Supervision, L.W., X.Z.; Funding Acquisition, L.W., X.Z., S.P.

## Supplementary Information

**Table S1.** Parameters (mean ± SD) of the fixations in the behavioral task: number of fixations, duration of fixations, dispersion of fixations (distance to the screen center).

|  | Face |  | House |  |
| --- | --- | --- | --- | --- |
|  | mean | SD | mean | SD |
| Number of Fixations | 3.8 | 0.9 | 3.7 | 0.9 |
| Duration of Fixations (ms) | 268 | 53 | 273 | 53 |
| Dispersion of Fixations (°) | 3.0 | 3.2 | 3.1 | 3.3 |

### Behavioral results

In the behavioral task, an N-back task was introduced to engage the picture viewing, which showed a 85.2 ± 11.1% (mean ± SD) hit rate, and a 4.7 ± 5.0% false alarm rate. In the Gaze Session, participants were informed to detect a subtle change in 20% of the trials (e.g., a flash of a black dot inside the moving dot), which showed a 90.0 ± 1.4% hit rate and a 0.2 ± 0.3% false alarm rate. In the Image Session, a 1-back task was introduced and all participants showed a hit rate above 95% with no false alarms.

### Cross-experiment classification of gaze patterns

To investigate if the distinct gaze patterns recorded in the MEG were related to the distinction between face and house categories, we performed cross-experiment classification where the classifier was trained with the gaze patterns from the behavioral experiment and was used to predict the gaze patterns in the Gaze Session of the MEG experiment (SF vs. SH, OF vs. OH). Moreover, to evaluate the sensitivity of self-generated gaze tracks, the classification analysis was performed by varying the number of fixations included. The cross-experiment classifications showed above-chance accuracies regardless of the included number of fixations, all *p* < 10^5^ except p = 0.011 for OF vs. OH when only the first fixation was included (Bonferroni-corrected, Fig. 1e in the main text). Importantly, the accuracies increased when the included number of fixations increased from one to two, but did not further increase when there were more than two fixations, all *p* < 0.001 for Fix 1 vs. Fix 1-2, Fix 1 vs. Fix 1-3, and Fix 1 vs. Fix 1-4, *p* = 0.1 for Fix 1-2 vs. Fix 1-3, *p* = 0.004 for Fix 1-2 vs. Fix 1-4, *p* > 0.999 for Fix 1-3 vs. Fix 1-4 (Bonferroni-corrected, Fig. 1e in the main text). Moreover, the accuracy for SF vs. SH was higher than the accuracy for OF vs. OH, but the difference held only when the first two fixations (Fix 1: *p* = 0.008, Fix 1-2: *p* = 0.032), not when more fixations were included (Fix 1-3, *p* = 0.088, Fix 1-4: *p* = 0.34, Bonferroni-corrected, Fig. 1e in the main text).

### The structural pattern of face-related gaze tracks

We have assumed that face-related gaze tracks had a structural pattern than house-related gaze tracks, hence treating the house-related gaze tracks as a control in the analysis of MEG signals. As an assumption check, we quantified the structural patterns of the gaze tracks by calculating the representational distance among the gaze tracks in different trials, assuming that a lower representational distance indicates a higher representational similarity^1^, hence a higher structured pattern^2^. The Euclidean distance was calculated based on the fixation parameters within the face category, within the house category, and between the two categories in the Gaze Session. The representational distance for face-related gaze tacks was lower than the representational distance for house-related gaze tracks, all *p* < 0.001 (Bonferroni-corrected), and also lower than the between-category representational distance, all *p* < 0.01, regardless of whether the gaze tracks were from the current observer (self-gaze, Fig. S1 left) or from another observer (other-gaze, Fig. S1 right). However, the representational distance for house-related gaze tracks was not significantly different from the between-category representational distance, all *p* > 0.38. This pattern still held even when more fixations were considered (Fig. S1). Consistent with Wang et al^2^, these results showed that the face-related gaze tracks had a highly structured pattern, whereas the house-related gaze tracks did not have such a consistent structure.

---

**Figure S1.**
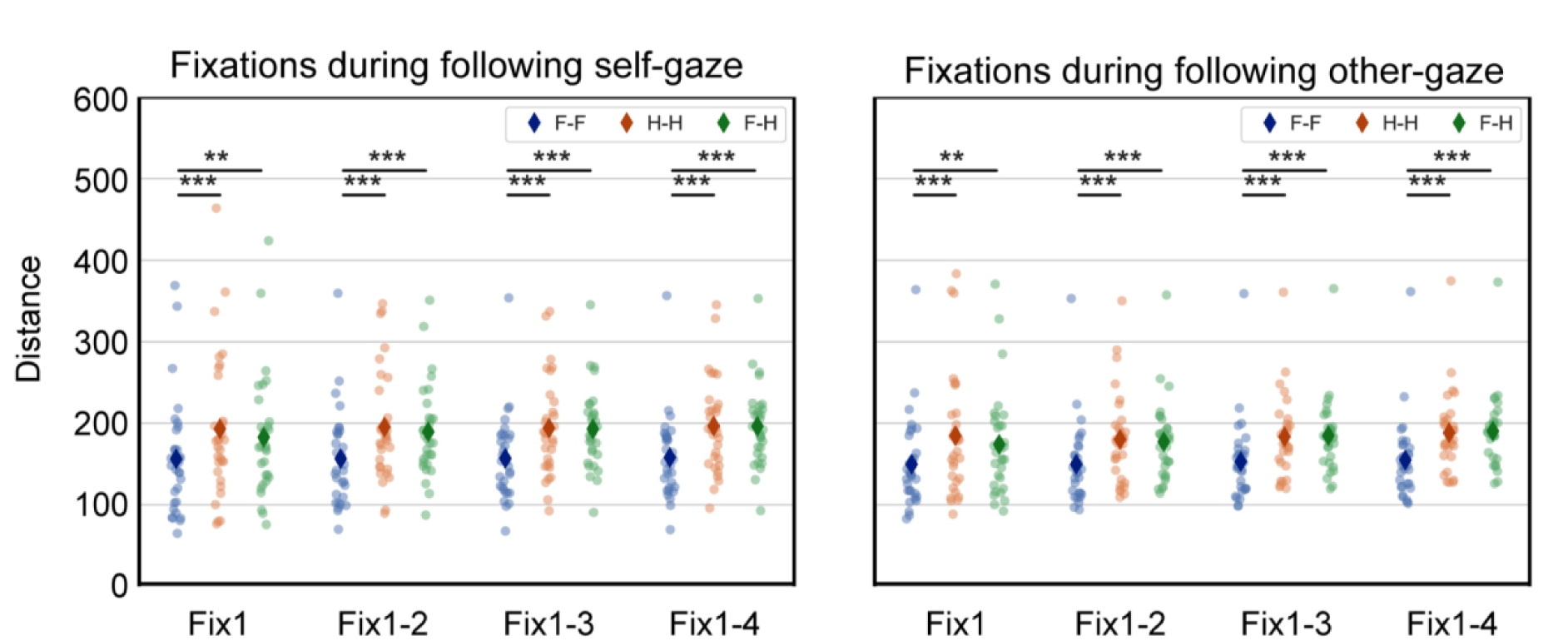
Structural pattern of face-related gaze tracks. The representational distance between fixations during the following of self-gaze tracks (left) and other-gaze tracks (right). F-F: the representational distance of fixations within the face category. H-H: the representational distance of fixations within the house category. F-H: the representational distance of fixations between face and house. The number of fixations was also included to show a consistent pattern. *** *p* < 0.001, ** *p* < 0.01.

**Figure S2.**
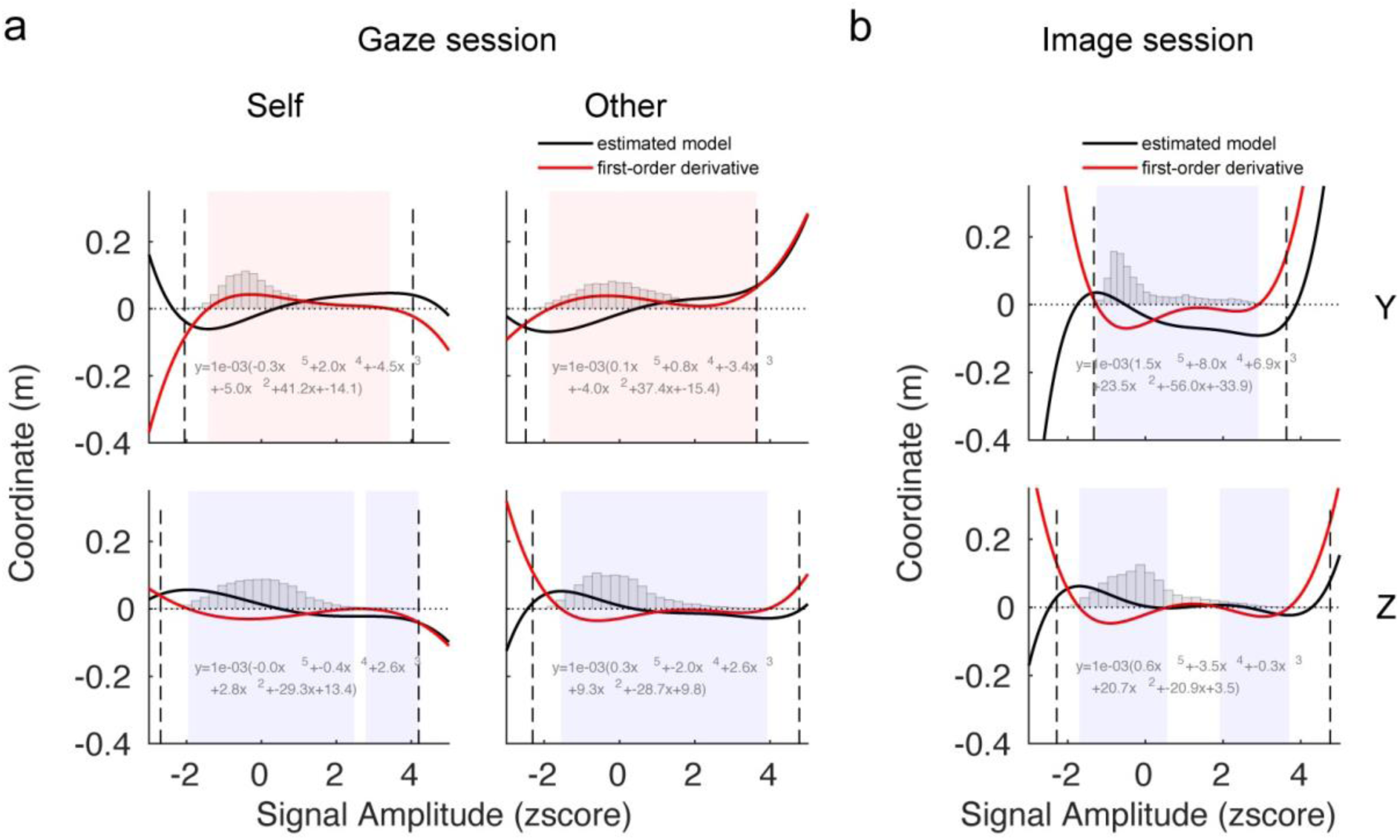
The estimated models (*y* dimension in upper row, *z* dimension in lower row) and their first-order derivatives of the MEG spatial gradients in the gaze session (**a**, left: SF vs. SH, right: OF vs. OH) and the Image Session (**b**, Face vs. House). The dashed vertical lines denote the minimum and maximum MEG signals, and the shaded areas denote the range that the derivatives > 0 (pink) or < 0 (violet). The gray histograms illustrate the distribution of the signal amplitudes.

